# *NUCOME*: A Comprehensive Database of Nucleosome Organizations in Mammalian Genomes

**DOI:** 10.1101/414094

**Authors:** Xiaolan Chen, Hui Yang, Yong Zhang

## Abstract

Nucleosome organization is involved in many regulatory activities in various organisms. However, studies integrating nucleosome organizations in mammalian genomes are very limited mainly due to the lack of comprehensive data management. Here, we present *NUCOME*, which is the first database to organize publicly available MNase-seq data resource and manage unified processed datasets covering various cell types in human and mouse. The *NUCOME* provides standard, qualified and informative nucleosome organization data at the genome scale and at any genomic regions for users’ downstream analyses. *NUCOME* is freely available at http://compbio.tongji.edu.cn/NUCOME/.

## Introduction

The nucleosome is the fundamental unit of eukaryotic chromatin and is involved in regulatory activities through interactions with DNA-binding proteins, including regulatory factors, chromatin remodelers, histone chaperones and polymerases (1). Genome-wide nucleosome organization maps have been established in multiple species, and some consistent nucleosome positioning patterns of specific regulatory elements have been reported. For example, nucleosomes act as barriers to transcription factors (TFs) interacting with *cis*-regulatory elements. Nucleosome-free regions (NFR) and regularly spaced nucleosome arrays are strongly associated with transcription initiation (1-4). Nucleosome remodelers are essential for nucleosome dynamics that remove, slide and reposition nucleosomes to overcome barriers and facilitate transcription initiation and elongation (3,5-7). The conserved nucleosome positioning patterns on transcription start sites (TSSs) and other regulatory elements highlight the importance of the nucleosome in regulatory activities.

Furthermore, previous studies have shown that tissue- and disease-specific nucleosome organization widely exists in the mammalian genome and is involved in cell differentiation (8,9), reprogramming (9,10), tissue impairment (11-13) and diseases (14-16). Most studies focus on identifying specific nucleosome organization features and associating these features to gene transcription or chromatin modifications. Cell type-specific nucleosome organizations typically indicate the chromatin environment that involves distinct regulatory factors and cellular processes (17). Therefore, nucleosome organization maps are critical for deriving a panoramic view regarding the chromatin structure, modification and their relationship in regulatory function. However, compared to other types of regulatory landscapes, such as histone modification, DNA methylation and transcription factors, nucleosome organizations have not been sufficiently explored.

To systematically explore the regulatory function of the nucleosome, a comprehensive and dedicated database of nucleosome landscapes is urgently needed to manage, explore and analyze these data resources. MNase-seq is the most widely used technology for generating nucleosome organization maps (18,19). The large size of mammalian genomes is a major challenge in experiments that require a very high sequencing depth, thus, MNase-seq data analyses are complicated and time-consuming. We established a comprehensive database named *NUCOME* (<u>Nuc</u>leosome <u>O</u>rganizations in <u>M</u>ammalian G<u>e</u>nomes, Figure 1) that organizes extensive MNase-seq data and characterizes nucleosome organizations. The following are the three major features of *NUCOME* database: 1) *NUCOME* is one of the most extensive catalog of MNase-seq data and is managed via a standard analysis pipeline and quality control (QC) metrics. 2) *NUCOME* provides high-quality nucleosome organization information for various human and mouse cell and tissue types. 3) *NUCOME* provides a web interface containing multiple modules, including data search, nucleosome organization visualization and an application that quantifies informative nucleosome organization data in genomic regions.

**Figure 1.**
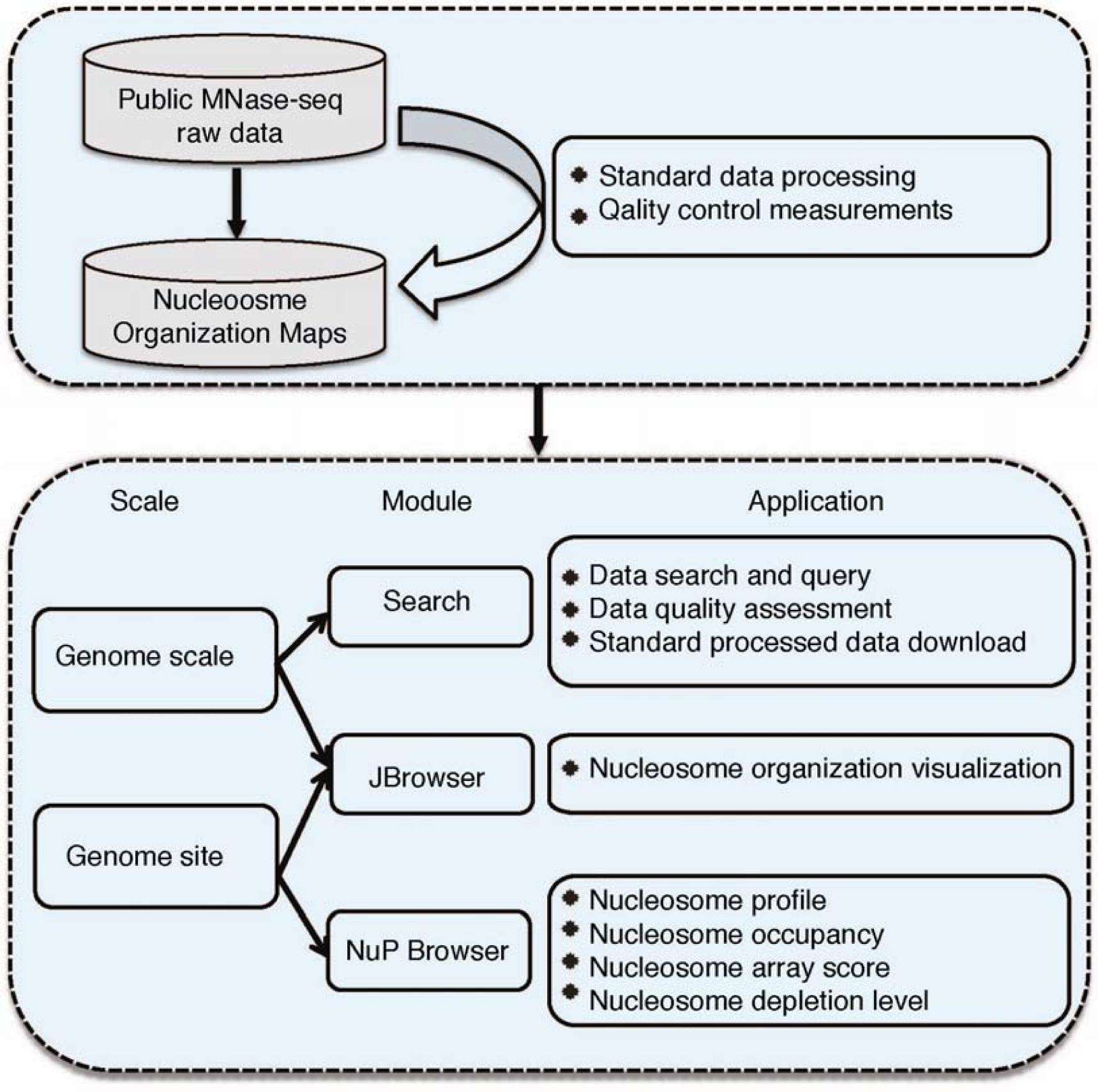
Structure of the *NUCOME* database.

## MATERIAL AND METHODS

### Calculation of QC measurements

NUCOME provides seven QC measurements for MNase-seq data to help users judge the data quality. The definitions of sequencing coverage, AA/TT/AT di-nucleotide frequency, nucleosomal DNA length, nucleosome depletion at TSSs, enrichment of well-positioned nucleosome arrays at DNase hypersensitive sites (DHSs) and untranslated regions (UTRs) were described previously (20). Nucleosome fuzziness downstream TSSs was defined as the coefficient of variance (CV) of the distance between +1, +2, +3 and +4 nucleosomes, while the position of nucleosomes was defined as the local maximum positions. For the measurement of nucleosomal DNA length, samples were ranked by the deviation of the measurement values to 146 bp in ascending order. For the measurement of nucleosome fuzziness downstream TSSs, samples were ranked based on the measurement values in ascending order. For each of other five measurements, samples were ranked based on the measurement values in descending order. For each sample, a “Pass” or “Fail” label was assigned for each QC measurement, except for sequencing coverage. The criteria of “Pass” and “Fail” for AA/TT/AT di-nucleotide frequency, nucleosomal DNA length, nucleosome depletion at TSSs, enrichment of well-positioned nucleosome arrays at DHSs and UTRs were defined previously (20). For nucleosome fuzziness downstream TSSs, samples with values lower than 0.4 was defined as “Pass”, while others were defined as “Fail”.

For each cell or tissue type, the reference nucleosome organization map was selected via two indicators. Firstly, the total number of ‘Pass’ QC measurements for each sample was defined as the first rank indicator. Then, we calculated the sum of rank quantiles of all QC measurements for each sample as the second indicators. For each cell or tissue type, the sample with the top rank for both two indicators was selected as the reference nucleosome organization map.

### Prediction of TF binding sites

We collected TF ChIP-seq data from Cistrome DB database (21) as the actual TF binding profile. Qualified ChIP-seq samples were selected by the QC measurements that provided in the database. For each TF ChIP-seq sample, the top 5,000 binding peaks (ordered by the fold change of peaks) were defined as the positive binding sites (bound sites). The negative binding sites (unbound sites) were defined as 20,000 random DHSs that do not overlap with any detected peaks in the ChIP-seq sample. All 25,000 bound and unbound sites were used for evaluating the TF binding prediction. Motif score was the first predictor. We scanned the entire genome for significant motif hits for each TF by using BINOCh (22). For any TF bound or unbound site with one or more significant motif hits, the highest motif matching score among those hits was assigned as the motif score of that site. For sites without any significant motif hits, their motif scores were assigned as 0. Nucleosome depletion level acted as the second predictor to improve the prediction performance. Here, the nucleosome depletion level at a given site was calculated as the degree of nucleosome occupancy deficiency at the center of the site. The maximum nucleosome occupancy at the centeral 200 bp bin of the site was defined as *N*_*cente*r_, and the maximum nucleosome occupancy surrounding the centeral bin spanning 200 bp on both sides was defined as *N*_*background*_. If *N*_*center*_ was larger than *N*_*background*_, the nucleosome depletion level of the site was assigned to 0. Otherwise, the nucleosome depletion level of the site was calculated as 1 – *N*_*center*_/*N*_*background*_.

A logistic linear regression conducted by a ‘glm’ function in R was performed to match the actual TF binding status (bound or unbound) by scoring motif score and nucleosome depletion level. The R package ‘pROC’ was used to evaluate the prediction power by calculating the true-positive rate and true-negative rate with different thresholds, and area under the curve (AUC) scores calculated by ‘auc’ function in R were used as an indicator to evaluate the performance of the TF binding prediction. The prediction improvement was calculated as AUC (motif score + nucleosome depletion level) – AUC (motif score).

## DATABASE CONSTRUCTION AND CONTENT

### *NUCOME* organizes extensive MNase-seq data of the mammalian genome

NUCOME focuses on human and mouse MNase-seq data. We collected and obtained 354 available data sets from Gene Expression Omnibus (GEO; https://www.ncbi.nlm.nih.gov/geo/) and downloaded raw data sets from Sequence Read Archive (SRA; http://www.ncbi.nlm.nih.gov/sra). The database comprises 28 mouse cell and tissue types and 25 human cell and tissue types (Table S1), representing one of the most extensive data sets of nucleosome organization in mammalian genome reported to date.

### *NUCOME* provides standard qualified nucleosome organization maps

To avoid variation in the processed data among the datasets, we applied a streamlined analysis pipeline starting with the raw sequenced data. We developed a standard workflow for managing MNase-seq data named CAM that includes read mapping, nucleosome organization profiling, nucleosome array detection an QC assessment (20). The workflow guarantees the quality of the data in the database and offers a convenient approach for users to download processed, standard and qualified data for their downstream analyses.

### Reference nucleosome organization maps selected based on quality comparison

To avoid users’ confusion in selecting high quality data among various samples, we determined reference nucleosome organization maps based on a quality comparison. We compared the data quality of samples from each cell and tissue type based on MNase-seq data specific QC measurements. The following seven QC measurements were introduced: sequencing coverage, AA/TT/AT di-nucleotide frequency, nucleosomal DNA length, nucleosome depletion at TSSs, nucleosome fuzziness downstream TSSs, enrichment of well-positioned nucleosome arrays at DHSs and UTRs (see Methods for details). We ranked the samples from the same cell or tissue type based on the values of QC measurements, and the top sample of each cell or tissue type was selected as the reference nucleosome organization map.

We integrated standard processed data for all samples and the reference nucleosome organization maps in a publicly accessible database named *NUCOME* (http://compbio.tongji.edu.cn/NUCOME/). The standard processed data for all samples allow users to conduct their analyses without difficulties in data pre-processing and QC. The reference nucleosome organization maps provide users with direct, quantified and reliable nucleosome organization information for any genomic region. *NUCOME* also retains the information of experimental condition for advanced users who are experts in nucleosome field to select suitable datasets for their analyses.

## RESULTS AND DISCUSSION

NUCOME includes three main modules (Figure 1), i.e. (a) ‘Search’, (b) ‘JBrowser’ and (c) ‘NuP Browser’ (nucleosome positioning browser), and the usage and interpretation of each module is briefly explained below. We present two examples of the usage of *NUCOME*. The first example involves a traditional analysis of MNase-seq data by aggregating nucleosome profiles around TSSs to illustrate the association of nucleosome organization and transcription activity. The second example focuses on the expanded usage of MNase-seq data for predicting TF binding events to illustrate interaction mechanisms of nucleosome organizations with other regulatory factors.

### Quality-supervised data query guides users in filtering samples

The ‘Search’ module in *NUCOME* allows for data retrieval in a quality-supervised manner to help users select data (Figure 2a). Two layers of QC assessments are included. The first layer displays the QC status of each QC measurement to allow users to judge the data quality directly (Figure 2a middle, Supplementary Figure 1a, b). In addition, users can compare samples from the same cell or tissue type in terms of each QC measurement using a polar plot (Figure 2a bottom, Supplementary Figure 1c, d) to filter and select high quality samples. The black line represents the average rank quantile of the QC scores for all samples in the corresponding cell or tissue type, while the blue line represents the rank quantile of the selected data. Supplementary Figure 1c presents an example in which the data quality of the sample is better than the average quality of all samples from the same cell or tissue type. In contrast, Supplementary Figure 1d presents an example of the sample with relatively worse data quality.

**Figure 2.**
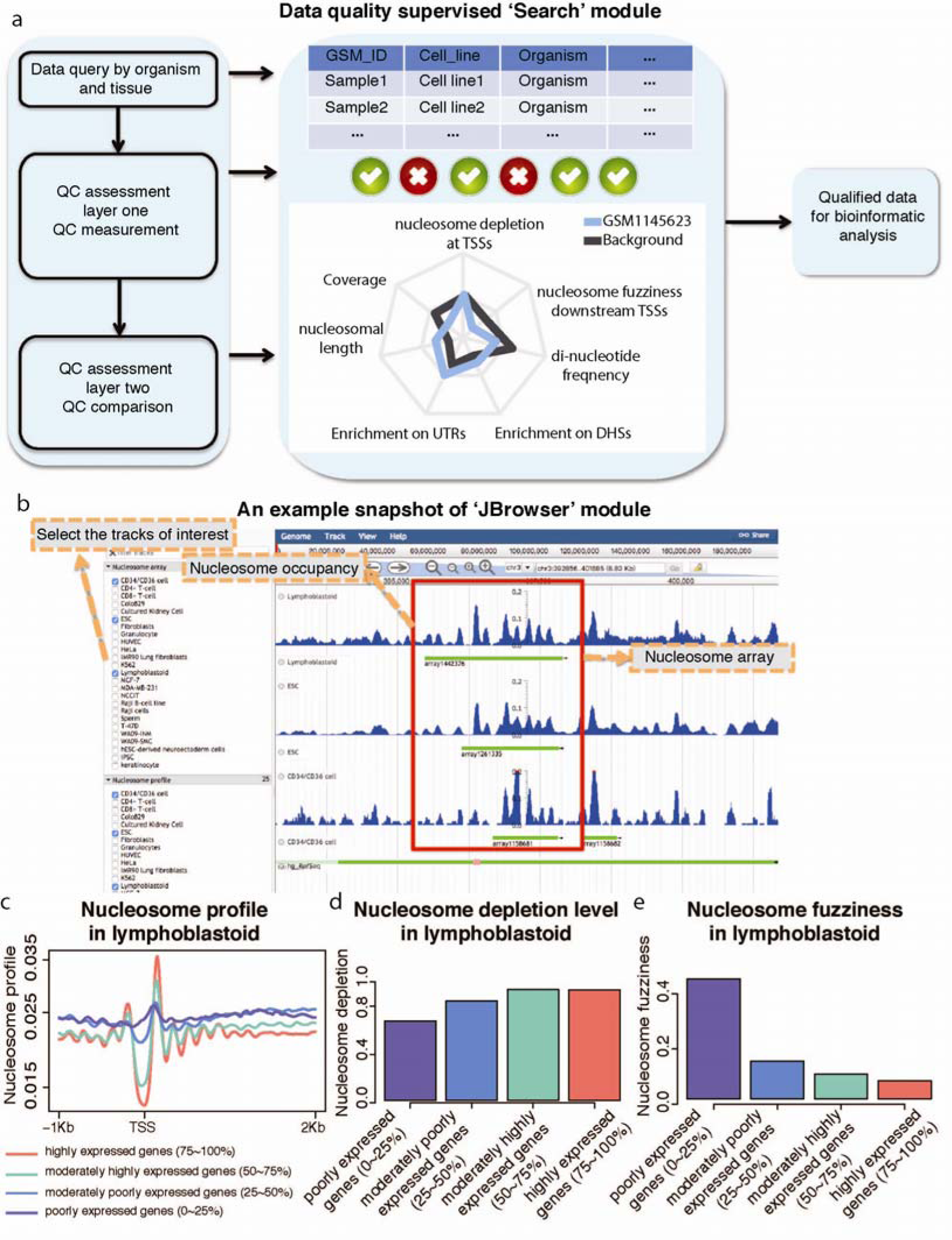
Examples of database usage. **a**) *NUCOME* provides a data quality supervised ‘Search’ module. Two layers of QC assessments are included. The first layer assesses the data quality of each sample as ‘Pass’ or ‘Fail’ for each QC measurement. The second layer QC assessment compares the data quality among samples from the same cell or tissue type by the rank quantile of the measurement value of each QC measurement. The black line in the polar plot represents the average rank quantile of the QC scores for all samples in the corresponding cell or tissue type, while the blue line represents the rank quantile of the selected data. **b**) An example snapshot of the usage of the ‘JBrowser’ module. The red box shows the location of nucleosome arrays in three tissues. **c**) The average nucleosome profiles around TSSs of four gene groups classified by expression level. **d**) Nucleosome depletion level around TSS of four gene groups classified by expression level. **e**) Nucleosome fuzziness downstream TSS of four gene groups classified by expression level.

In addition, the ‘Search’ module provides other useful information about the samples. The ‘sample description’ section includes brief and basic information about the samples. The ‘nucleosome array’ section presents the locations of detected nucleosome arrays around TSSs ordered by the enrichment of the arrays on the promoters. The ‘download’ section provides access to processed files, including mapped reads, nucleosome occupancy and nucleosome arrays.

### Data visualization of reference nucleosome organization maps

The ‘JBrowser’ module provides an overview of the reference nucleosome organization maps of the entire genome queried by gene or genomic region (Figure 2b). The nucleosome organization maps focus on genome-wide nucleosome occupancy and the location of nucleosome arrays. Nucleosome occupancy is measured as the number of reads mapped to the genomic site in a cell population and presents the probability that a nucleosome is localized at the genome site. Thus, nucleosome occupancy is an important feature that reflects DNA accessibility. It has been reported that well-positioned nucleosome arrays downstream TSSs participate in transcription regulation with either positive or negative effects (1) in a cooperative context with other regulatory factors (23). We hypothesized that the location of nucleosome arrays may indicate significant regulatory activities. Thus, we display the nucleosome occupancy and nucleosome arrays in the module of data visualization to present nucleosome organization on a genome-wide scale.

### Informative nucleosome organization illustrating regulatory function

The ‘NuP Browser’ module provides users with a friendly and flexible platform to query nucleosome organization information in any genomic region or any gene. Compared with the ‘JBrowser’ module, ‘NuP Browser’ module provides more nucleosome organization features and quantifies the information as easily processed scores describing the local chromatin structure, including nucleosome occupancy, nucleosome array score, nucleosome depletion level and nucleosome profile. The text format output allows users to perform nucleosome positioning analyses without encountering difficulties in data processing.

The nucleosome typically participates in biological processes by interacting with other regulatory factors. Here, we used an example to explore the function of nucleosome organizations around TSSs in transcription activity. We profiled aggregated nucleosome positioning patterns around TSSs of genes with different expression levels and compared the differences between the nucleosome positioning patterns (Supplementary Figure 2a). The analysis illuminated the relationship between nucleosome organization and transcription activity. The promoters of highly expressed genes tend to exhibit stronger canonical nucleosome organization, including nucleosome depletion level at TSSs and a regularly spaced nucleosome array downstream the TSSs (Figure 2c). The better nucleosome positioning pattern exhibits an increased nucleosome depletion level (Figure 2d) and reduced nucleosome fuzziness (Figure 2e). Many previous studies have reported similar observations using specific cell cultured systems or tissues(24-30). Here, we showed that the regulation manner is conserved among various human and mouse tissues and cell types by performing this analysis in multiple samples stored in *NUCOME* (Supplementary Figure 2b-e). Using this comprehensive database, researchers can deeply explore the features of nucleosome-mediated regulation.

### Nucleosome organization influences transcription factor binding

Previous studies have confirmed that nucleosome positioning patterns around TFs may indicate the regulatory function of the TFs and play a predictive role in gene expression (31). Here, we explored the ability to predict TF binding *in vivo* by introducing informative nucleosome organization (Supplementary Figure 3a). We used a logistic linear regression to match the binding status of TFs in 140 ChIP-seq datasets of TFs covering 7 different cell types and 73 TFs in human by scoring motif score of DNA sequencing and nucleosome depletion level for those sites. Then, we applied a Receiver Operating Characteristic (ROC) curve analysis to evaluate the performance of the prediction based on AUC scores (see Methods for details). The results demonstrate that the AUC scores improve universally after including the nucleosome depletion level in the prediction model (Figure 3a). The improvements may result from the effect of nucleosome organizations on chromatin accessibility for TFs binding. The improvements are not equal in all TFs, and the TFs can be classified as Improve_G and Improve_S groups, with great or slight improvement of prediction power by adding nucleosome depletion level (Figure 3b). We found that the prediction improvements in TF binding prediction are significantly negatively correlated with the intrinsic DNA motif prediction contribution (Figure 3c). Next, we divided TFs into two groups, Motif_H and Motif_L, with higher or lower DNA motif prediction contribution separately (Figure 3d). The TFs in Improve_G and Motif_L groups are significantly overlapped (Figure 3e), indicating the valuable contribution of nucleosome organization in TF binding prediction, especially for TFs with poor prediction power based on DNA motifs. Similar results were also observed for TFs binding prediction in mouse (Supplementary Figure 3b, c). Taken together, these results demonstrate that nucleosome organization information derived from *NUCOME* can be widely applied to elucidate the transcription regulatory program in human and mouse.

**Figure 3.**
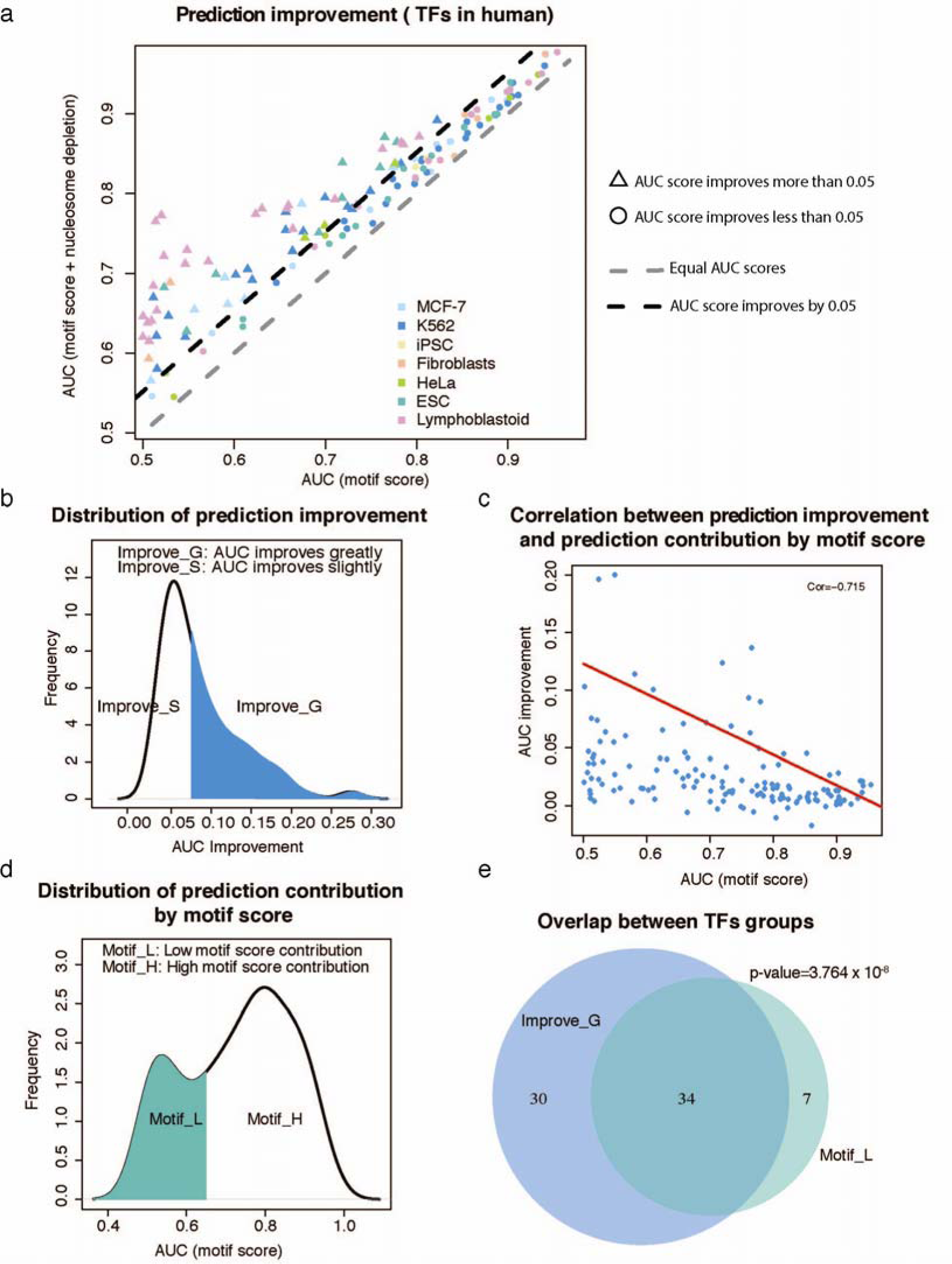
Nucleosome organization information improves TF binding prediction. **a**) A scatter plot displays the improvement of prediction power by introducing nucleosome depletion level in the prediction model. The x-axis represents the AUC score by using DNA motif score only. The y-axis represents the AUC score by using both DNA motif score and nucleosome depletion level. Different colors represent the TFs in various cell and tissue types. The gray dot line indicates that the AUC scores of the two predictions are equal. The black dot line indicates that adding the nucleosome depletion level improves the prediction power by 0.05. The triangle points are TFs that AUC scores improve more than 0.05, while the round points are TFs that AUC scores improve less than 0.05. **b**) The distribution of AUC score improvements by introducing nucleosome depletion level in the prediction model. ‘Improve_G’ group includes TFs exhibits at least a 0.05 improvement, while ‘Improve_S’ group includes TFs with less than 0.05 AUC score improvement. **c**) A scatter plot with the red regression line revealing the significantly negative correlation between prediction improvement by introducing nucleosome depletion level and prediction performance by DNA motif score only (p-value < 2.2 x 10^-16^). The x-axis represents the AUC score by using DNA motif score only. The y-axis represents the AUC score improvements by introducing nucleosome depletion level. **d**) The distribution of prediction performance by DNA motif score only. ‘Motif_L’ group includes TFs that have AUC scores less than 0.65 by using DNA motif score, while ‘Motif_H’ group includes TFs with AUC scores higher than 0.65. **e**) Overlapping among TFs in ‘Improve_S’ group and ‘Motif_L’ group. The p-value is calculated by the Chi-square test.

## CONCLUSIONS

*NUCOME* is a comprehensive database that organizes the most extensive data sources of MNase-seq data and provides standard nucleosome organization maps of various cell and tissue types in human and mouse. *NUCOME* provides three modules in a web interface for 1) querying data, 2) visualizing reference nucleosome organization maps and 3) querying nucleosome organization information in any genomic regions and providing text format outputs for users’ downstream analyses. Given that nucleosome organization participates in various regulatory activities, and *NUCOME* is the first comprehensive database of nucleosome organization data, the database can be a valuable resource to elucidate the panoramic view of transcription regulatory program in human and mouse.

## AVAILABILITY

The database is available at http://compbio.tongji.edu.cn/NUCOME/.

## Acknowledgements

We thank Ji Liao for his contribution in the early stage of this project.

